# Within-Subject Precision Measurement: Theory and Utility of Assessing Change Scores and Their Variance

**DOI:** 10.64898/2026.11.16.767165

**Authors:** Randy L. Buckner, Deanna M. Barch, Patrick Mair, Thomas E. Nichols, Theodore D. Satterthwaite, B.T. Thomas Yeo

## Abstract

Within-subject studies gain statistical power by removing stable between-subject differences from the comparison. However, the metrics commonly used to select measurement instruments and anticipate their within-subject performance are often based on between-subject correlations of raw scores from test-retest designs. Measuring each subject’s change directly – after an intervention, and ideally more than once – provides a firmer basis for evaluating measurement precision and planning studies of within-subject change. Here we describe the statistical theory behind estimating measurement properties from change scores. In one instantiation, the pre- and post-intervention measurements are each made twice, yielding the duplicate change-score design. Such a design provides an estimate of the mean signal change and enables the decomposition of the total observed variance of change scores into true between-subject response heterogeneity and measurement noise. In extended designs, where multiple measures are collected within and across days, measurement noise can be further decomposed to distinguish within-session from between-day measurement variability. These estimates supply the inputs needed for within-subject study design, including providing a basis for selecting among alternative measurements and comparing processing pipelines, performing power calculations, and estimating the precision limits of detecting change within individual subjects.

---

Determining a measurement instrument’s sensitivity to detect within-person change is central to translational research and has grown more challenging as measurement options have multiplied. Rich data streams and flexible processing pipelines – spanning wearable sensors, neuroimaging, and language-model scoring – require a cascade of decisions before a change score can be formed (e.g., Carp, 2012; Botvinik-Nezer et al., 2020; Bhagwat et al., 2021; Schweinsberg et al., 2021; Rijnhart et al., 2022; Breznau et al., 2022). Each decision can affect both the signal and the noise.

From a power perspective, the task is to select a valid instrument and processing pipeline that minimize measurement error while preserving the signal of interest. The commonly used tool for evaluating instrument reliability is the intraclass correlation coefficient (ICC) computed on repeated measurements under the same conditions spaced close in time^1^ (Shrout and Fleiss, 1979; McGraw and Wong, 1996). For discriminating among individuals, this use of the ICC is appropriate: establishing normative ranks or predicting who will respond to treatment both benefit from a high ICC, which indicates that the instrument consistently rank-orders subjects.

A high ICC, however, does not answer the question that within-subject designs depend on: how precisely can change be measured within a subject? High ICC values occur when between-subject variation is large relative to measurement error. Thus, a large measurement error can be hidden by even larger between-subject variation, and a genuinely precise instrument with low measurement error can be undervalued simply because true between-subject variability is small (for discussion see Bland and Altman, 1986; Guyatt, Walter, and Norman, 1987; Mellenbergh and van den Brink, 1998; Haertel, 2006; de Vet et al., 2011). For clinical intervention trials in particular, the central question is whether an instrument can capture meaningful change in the construct being measured, a characteristic sometimes described as responsiveness (Mokkink et al., 2010; see also Guyatt, Walter, and Norman, 1987).

This has direct consequences for power analyses. Accurate sample-size calculations for within-subject designs require three main inputs: the measurement error of the instrument (measurement noise), the average expected change (the change signal), and the between-subject variability of that change (response heterogeneity). Traditional test-retest scores yield measures of within-subject measurement error – the standard error of measurement (SEM)^2^ and the related minimal detectable change (MDC) – both expressed on the original measurement scale and both useful for power analyses (Weir, 2005). These metrics are nonetheless blind to processing effects on the change signal.

Consider the three data scenarios shown in Figure 1. In each scenario, a set of within-subject change scores is illustrated. The scenarios differ in their level of between-subject baseline differences (i.e., true between-subject variance), measurement noise, and the recovered magnitude of the signal change. None of the quantitative measures typically used to assess reliability provide information about whether the measurement process (e.g., the type of processing pipeline used) attenuates the recovered signal. For example, the large signal differences between the first and third columns in Figure 1 are not revealed in the correlation plots (panels G and I), even though the change signal is markedly reduced.

**Figure 1.**
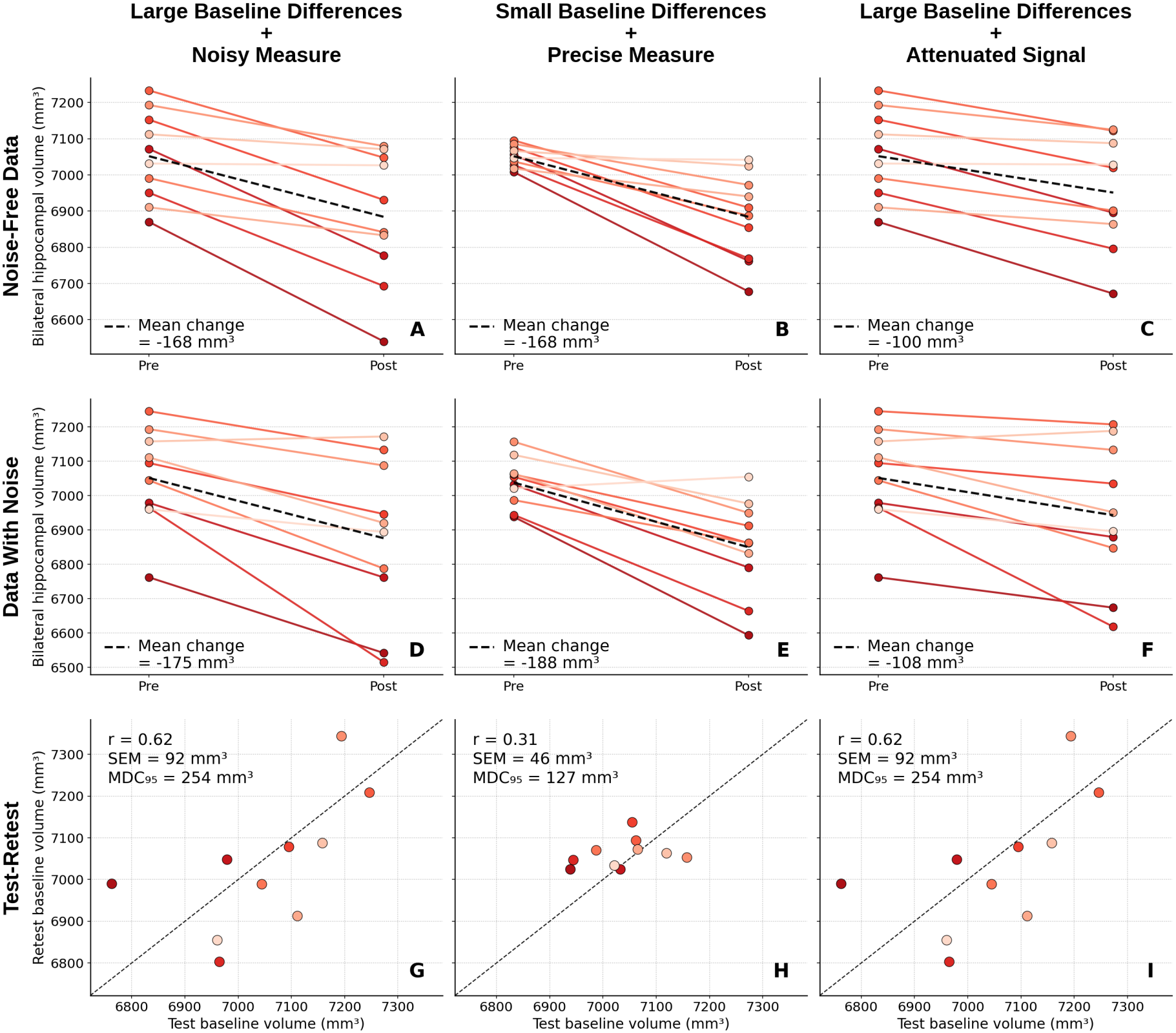
Ten subjects are simulated over a pre–post interval during which change in hippocampal volume (in mm^3^) is measured. Each line is a simulated subject. True response SD (*σ*_response_) is held constant, and individual deviations from the population change are color-coded from deep red (large decline) to pale (minimal decline). The **Large Baseline Differences + Noisy Measure** column (**A, D, G**) uses a cohort with large baseline differences between individuals (*σ*_baseline differences_ = 122 mm^3^) paired with a noisy instrument (*σ*_error_ = 80 mm^3^). The **Small Baseline Differences + Precise Measure** column (**B, E, H**) uses a less heterogeneous cohort (*σ*_baseline differences_ = 29 mm^3^) paired with a more precise instrument (*σ*_error_ = 40 mm^3^). **The Large Baseline Differences + Attenuated Signal** column (**C, F, I**) uses the same cohort as the first column while attenuating (reducing) each subject’s recovered response by 40% of its true magnitude. The **Noise-Free Data** row shows data from the three scenarios without measurement error. The **Data With Noise** row adds measurement noise. The **Test-Retest** row plots two independent baseline measurements from each scenario as is commonly obtained in a conventional test-retest reliability study. Note that the typically available test-retest metrics (correlation coefficient, r; standard error of measurement, SEM; minimal detectable change, MDC) do not fully characterize the sensitivity to detect change. For example, note that the r values in panels G and H differ in the direction opposite to the improvement in measurement error reflected by SEM and MDC. Raw-score SEM and MDC capture the measurement error but do not detect signal attenuation (panels G and I are indistinguishable even though panels A and C clearly differ). In this paper, we illustrate how replicate change scores can decompose sources of measurement error while simultaneously estimating the change signal, providing the full set of measurement properties needed for within-subject study design and power analysis.

This blind spot matters because acquisition and processing choices can affect the change signal in hard-to-predict ways, alongside their intended effect on noise. As researchers compare many processing alternatives, optimization may select candidates that trade signal attenuation for noise reduction in ways that reduce power to detect within-subject change. In fMRI, shortening the echo time can lower measurement error but, past a point, also weakens the functional signal (Stöcker et al., 2006). An overly large spatial-smoothing kernel can reduce noise while also attenuating the signal it is meant to recover (Mikl et al., 2008). As processing options multiply, a challenge becomes finding pipelines that reduce error while also ensuring that the measured signal is retained.

The present paper uses a duplicate change-score design to characterize the reliability of measurement instruments and processing streams. Within such a design, the baseline (pre-intervention) and follow-up (post-intervention) measurements are each repeated, so that the measurement error can be decomposed (Figures 2 and 3; Senn, 2018). Elliott et al. (2026) recently used this design to test the precision limits for detecting brain-structure change in individuals. The design separates two conceptually distinct quantities, the true response heterogeneity of the change (*σ*^2^_response_) and the measurement noise of within-individual change (*σ*^2^_error_). It also estimates the mean signal change 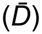, so that instrument and processing choices can be checked to determine whether they reduce (attenuate) the recovered signal. Because a within-subject paired contrast cancels individual trait-response heterogeneity, one can test effects on the signal with good power even when the absolute value of 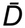 is uncertain. We develop this approach in the sections that follow, as well as suggest alternative change-score designs that can use fewer sessions.

**Figure 2.**
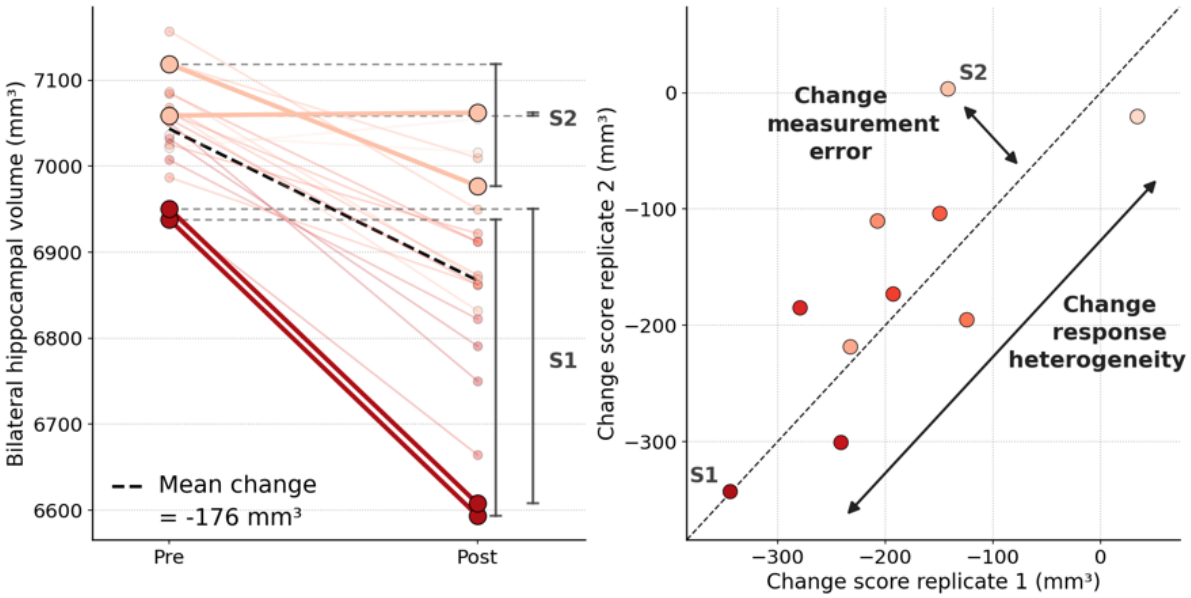
The duplicate change-score design allows measurement error to be separated from variability in the true response (signal). Left.The simulated subjects from Figure 1 (panel B) are replotted to highlight two subjects with each contributing replicate change scores (shown as the two lines for each). **S1** shows a large pre-to-post change response and **S2** shows a small response. Within the duplicate change-score design, two estimates (replicates) are obtained pre- and post-intervention providing two separate estimates that can be compared as illustrated in the right panel. **Right**. The replicate pre-to-post change scores are plotted for each subject color-coded to match their lines. When plotted in this manner, the response heterogeneity, which is reflected in each subject’s position along the X = Y identity line, is separated from the estimate of the measurement error, which is the distance from the identity line.

**Figure 3.**
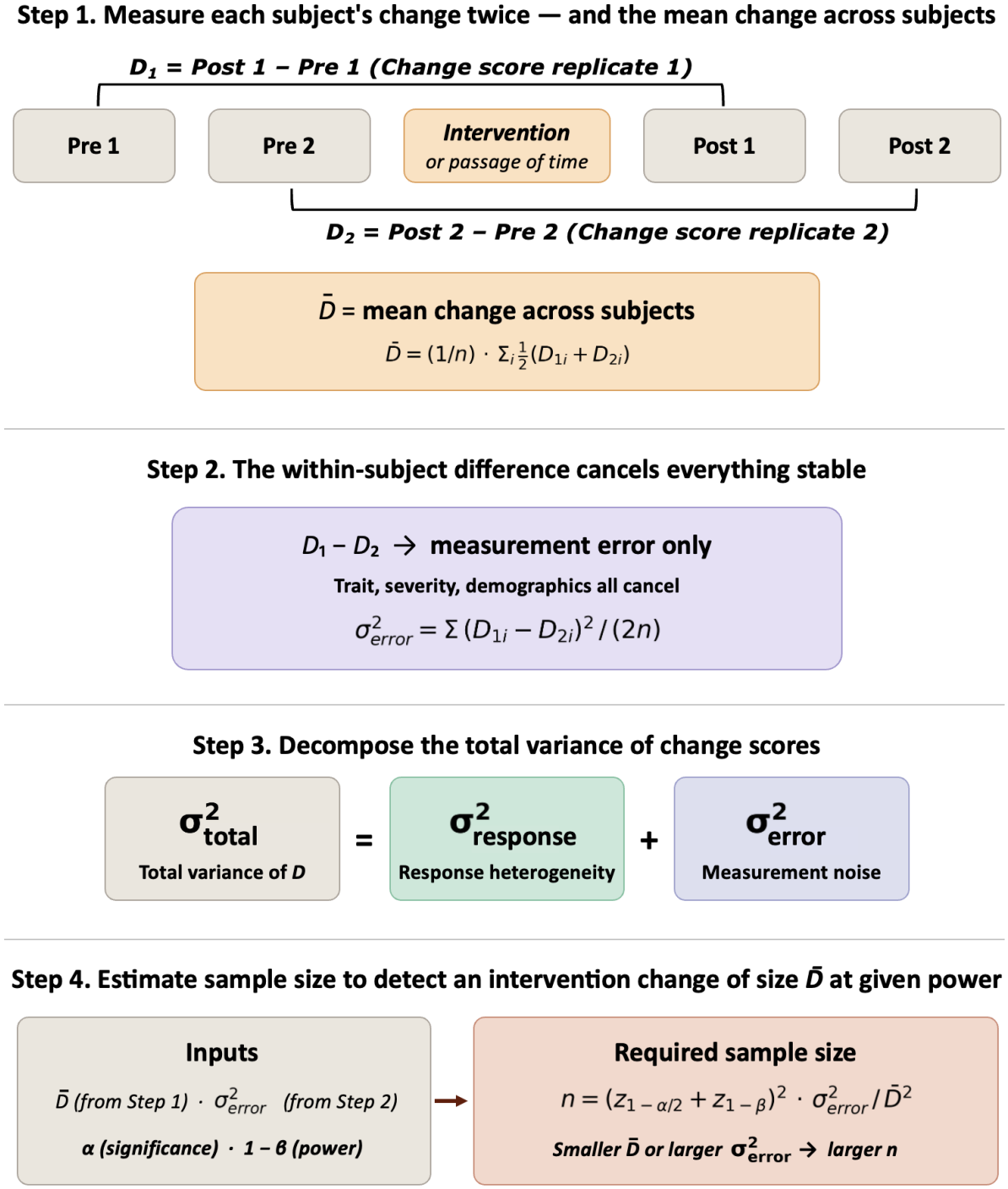
The duplicate change-score design provides a strategy to estimate the change signal and measurement error in a framework that supports power planning for within-subject research. Step 1.Each subject’s change score is measured twice in order to provide the information to decompose the sources of variance. Critically, the estimated change itself, 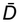, quantifies the signal. **Step 2**. The within-subject differences of the replicate change scores provide an estimate of the measurement error, independent of stable between-subject differences. **Step 3**. The total variance can be decomposed through subtraction to estimate the separate components due to between-subject response heterogeneity (*σ*^*2*^_response_), that is whether some subjects truly respond more than others, and measurement noise (*σ*^*2*^_error_), which reflects properties of the measurement instrument, including processing choices. **Step 4**. The three estimated quantities (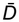, *σ*^2^_response_, and *σ*^2^_error_) are critical inputs for power planning and allow estimation of sample size for a contemplated within-subject research study.

Our approach draws on several methodological traditions (e.g., Cronbach et al., 1972; Bland and Altman, 1986; Guyatt et al., 1986; Senn, 2002, 2018). Statistically, the approach is a specific instance of a random-effects, generalizability-theory model applied to replicated change scores, and it can be handled within either frequentist or Bayesian frameworks. Our contribution is to link that theory to a concrete acquisition strategy anchored on change scores, in a form useful for comparing instruments and optimizing power in within-subject research, including the design of intervention trials.

## The Duplicate Change-Score Design

If within-subject design optimization requires preserving the mean signal change, 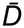, while minimizing the measurement noise of within-subject change, then a pilot or reliability study should be designed to estimate these directly, without the confounding influence of between-subject differences. This can be accomplished by measuring each subject’s change score twice around an intervention as illustrated in Figure 3.

Operationally, each subject in the reliability study completes the measurements on two separate occasions before and after the intervention, yielding two replicate estimates of change for each subject for the same intervention^3^. Letting *D*_*i*1_ and *D*_*i*2_ denote the two change scores for subject *i*, the within-subject difference *D*_*i*1_ − *D*_*i*2_ is free of any stable characteristic of subject *i* because quantities that are constant across the two occasions cancel out in the subtraction (e.g., baseline trait, severity, responsiveness, demographic characteristics). What remains is within-subject measurement noise, whose variance is estimated as follows:

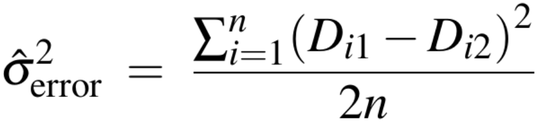

where *n* is the number of subjects in the sample, each contributing a pair of change scores (*D*_*i*1_, *D*_*i*2_). Here, 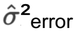 denotes the within-subject variance between replicates of the change score – that is, the average squared discrepancy between *D*_*i*1_ and *D*_*i*2_ within the same subject, scaled by 1/2 because the difference of two independent measurements has variance 2*σ*^2^_error_. Under the assumption of independent pre- and post-measurement errors^4^, *σ*^2^_error_ reflects the deviation of each subject’s measured change from their true change score value^5^.

This error quantity is informative for within-subject design power calculations but is often absent or neglected in study planning, although Bland and Altman (1986) and Weir (2005) describe similar quantities. It is derived without reference to between-subject variance and with minimal assumptions about the heterogeneity of the sample used to estimate it. A reliability study conducted in a homogeneous sample will likely yield estimates comparable to those from a heterogeneous sample, provided the measurement error structure is roughly the same. Estimates of measurement noise will nonetheless reflect characteristics of the sample from which they are obtained; non-compliant or otherwise inconsistent subjects will yield higher error estimates.^6^

## Variance Decomposition: Separating Measurement Error from True Response Heterogeneity

A key feature of the duplicate change-score design is that it enables separation of *σ*^2^_response_ (true between-subject response heterogeneity) from *σ*^2^_error_ (measurement noise). Under the model:

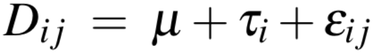

where *μ* is the mean change across subjects, *τ*_*i*_ is subject *i*’s true deviation from that mean, and *ε*_*ij*_ is the residual error of the change score for replicate *j* of subject *i*, so *D*_*ij*_ is the *j*-th replicate change score for subject *i*. The total variance is:

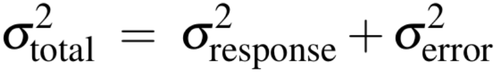

With *σ*^2^_total_ estimated from the observed change scores and *σ*^2^_error_ estimated from within-subject disagreement between *D*_*i1*_ and *D*_*i2*_, the between-subject response heterogeneity can be estimated by subtraction:

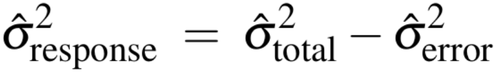

This decomposition has practical implications for study design. *σ*^2^_error_ reflects the within-subject residual variability of the change score (that is, measurement noise reflecting the precision of the instrument). It can often be reduced by averaging replicate measurements. For example, if each subject is measured twice at baseline and twice at follow-up, and the change score is formed from the averaged observations, the measurement noise contribution to the change score will be reduced. This represents a design-level adjustment that does not require modification of subject enrollment or the intervention itself.

By contrast, *σ*^2^_response_ reflects the true heterogeneity in how subjects respond to the intervention, independent of measurement noise. One subject might consistently have a large response to an intervention, while another might have a small response. Replicate averaging cannot reduce this between-subject response heterogeneity, as it may reflect real biological or psychological variability. It can, however, be addressed through stratification, covariate adjustment, or subject enrichment strategies that select individuals likely to show more uniform responses. Considering the magnitude of response heterogeneity (*σ*^2^_response_) relative to measurement noise (*σ*^2^_error_) might therefore help guide whether measurement refinement, design refinement, or both are likely to improve power.

The reliability of the change score (*ρ*_change_) can also be expressed as:

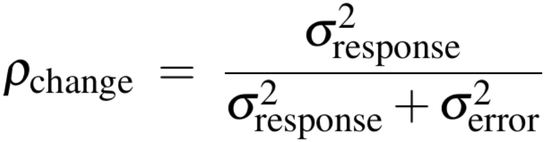

This quantity is a version of the classical ICC defined under a one-way random-effects model, but applied to change scores rather than raw scores. *ρ*_change_ isolates response heterogeneity (Figure 2; see also Senn, 2018). It answers the question: what proportion of the variation observed across individual change scores reflects real differences in response rather than measurement noise? When *σ*^2^_response_ is large relative to *σ*^2^_error_, individual response profiles are reliable, and response heterogeneity is a genuine feature of the intervention’s effect. When *σ*^2^_response_ is small, the primary determinant of observed change is measurement noise, and the instrument should be refined or averaged to maximize power.

## Power Analysis and Optimization Using the Duplicate Change-Score Design

The duplicate change-score design generates empirical inputs that can support power analysis for within-subject study designs, including future trials that themselves do not replicate measurements. That is, in many situations the duplicate change-score design, or a suitable simplified variant^3^, would be conducted in a pilot sample with the intent to use the estimates to design and power an intervention trial in which single pre- and post-measurements are collected. The test statistic (*t*) for a within-subject change (one pre-, one post-measurement) can be written as:

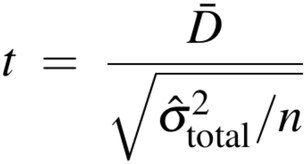

where 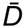 is the mean signal change, 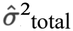 is the variance of change scores, and *n* is the number of subjects. For a two-sided test at significance level *α*, the required sample size (*n*) to achieve power 1 − *β* is:

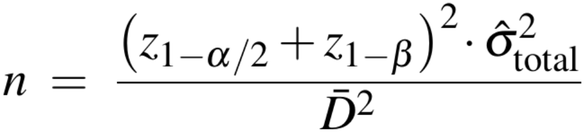

where z_1-*α*/2_ is the (1 − *α*/2) quantile of the standard normal distribution (the two-sided critical value at significance level α) and z_1-*β*_ is the (1 − *β*) quantile (corresponding to power 1 − *β*); for example, at α = 0.05 and 80% power, z_1-*α*/2_ ≈ 1.96 and z_1-*β*_ ≈ 0.84.

Following the variance decomposition described above, the duplicate change-score study yields:

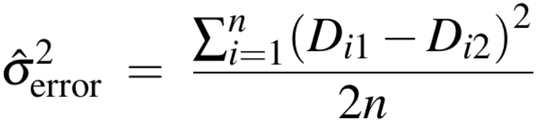

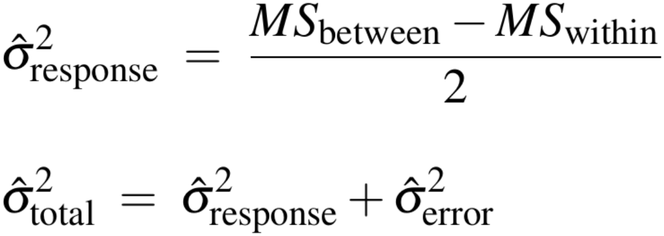

where MS_between_ and MS_within_ are the between- and within-subject mean squares from a one-way random-effects ANOVA on the 2*n* change scores, with subject as the random factor. Defined in this manner:

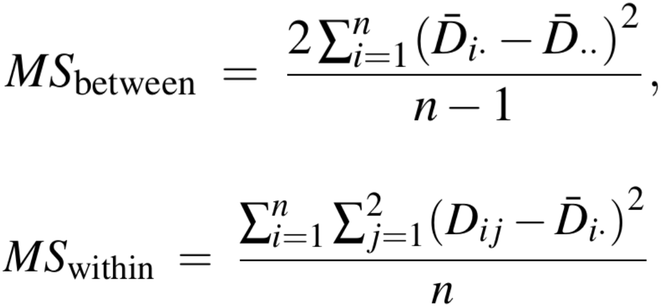

For the power analysis, the signal 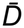 can be estimated from collected data as the grand mean of all 2*n* change scores across subjects and replicates (though be aware of the problems with retrospective power analysis [Thomas, 1997; Hoenig and Heisey, 2001; Wang, 2010]). Alternatively, it can be defined based on the smallest effect size of interest (SESOI) or minimal clinically important difference (MCID; e.g., Riesthuis, 2024). Together, these quantities parameterize the power calculation: 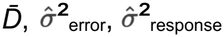 and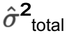. The estimate is grounded in the within-subject behavior of the instrument under an approximation of the target conditions. Power calculations can be based either on the mean change in natural units or on a standardized effect size such as Cohen’s *d*_*z*_. The effect size for the within-subject design expressed as Cohen’s *d*_*z*_ is:

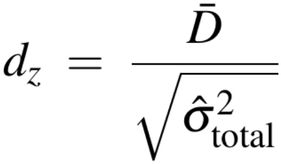

Because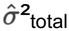 is derived from a design that isolates its two components, the investigator can inspect the relative contributions of 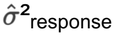 and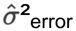 to determine whether the denominator is driven primarily by genuine response heterogeneity or by measurement noise.

## The Limits of Precision

The decomposition of 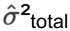 into 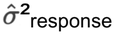 and 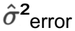 allows an investigator to estimate the limits of precision for both the measurement instrument and the individual response. These estimates have direct implications for how repeated measurements can be deployed in study designs to reduce sample size and, critically, reduce the MDC within an individual.

If *k* replicate measurements are obtained at each timepoint (at baseline and follow-up) and averaged before computing the change score, the effective error variance is reduced to 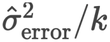, while 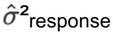 remains unchanged, because true response heterogeneity is a property of the cohort and intervention rather than the measurement process^6^. The modified power calculation becomes:

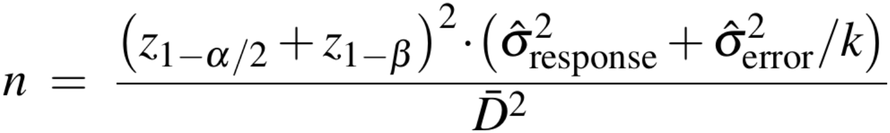

Under an ideal scenario, in which replicate measurements are independent, the gain in sample size efficiency from averaging k replicates per timepoint is:

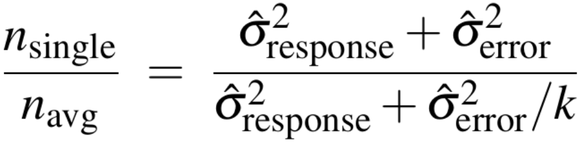

This ratio is large when 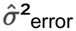 dominates and approaches unity as 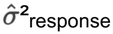comes to dominate. The duplicate change-score design thus enables the investigator to determine whether obtaining replicate measurements at each timepoint is justified by the reduction in required sample size.

The relation can be expressed as a power curve over k, with the asymptote representing the power achievable if measurement noise 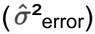 were eliminated entirely – a ceiling then determined by 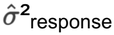. When this ceiling exceeds the target power, modest averaging of k = 2 to 3 replicates per timepoint may improve power. When 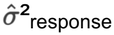 is itself large relative to the expected signal 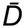, averaging offers diminishing returns, and enrichment strategies or covariate adjustment will reduce true between-person response heterogeneity. Figure 4 illustrates these points.

**Figure 4.**
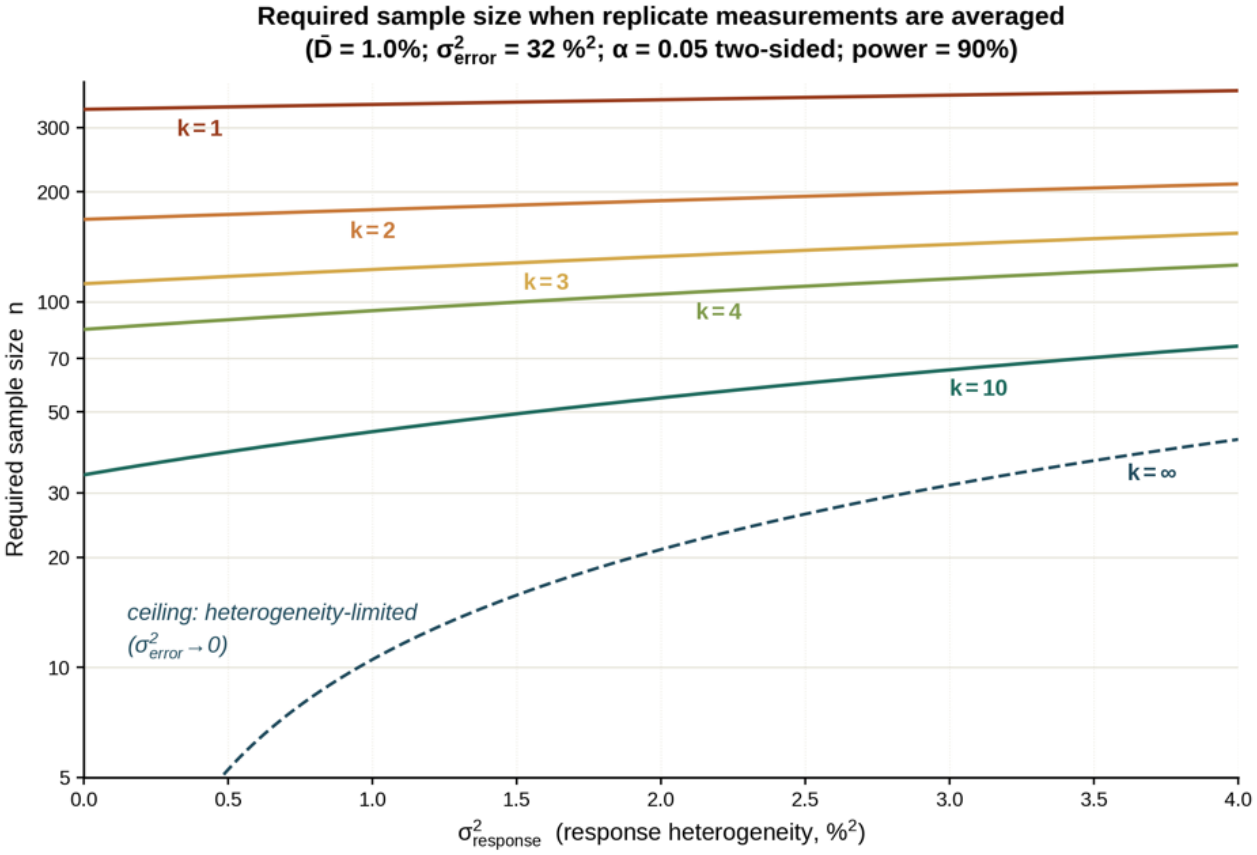
Required sample size to detect a change as a function of response heterogeneity (σ^2^_response_) when k replicate measurements are averaged at each timepoint, for k = 1 to 4, 10, and the ∞ ceiling. Averaging reduces the effective error variance to σ^2^_error_/k while leaving σ^2^_response_ unchanged. Power calculations assume 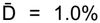, σ^2^_error_ = 32%^2^ (σ_error_ = 5.66%), α = 0.05 two-sided, and 90% power. The k = ∞ curve marks the heterogeneity-limited ceiling: as σ^2^_error_ → 0, required n depends only on σ^2^_response_, defining the ceiling on what replicate averaging alone can achieve. Note that required sample size n differs across the entire range of response heterogeneity (σ^2^_response_) reducing the sample size burden from n > 300 to n ≈ 100 when k = 4 replicates are obtained. In many situations the additional replicates are far less costly than the increased sample size.

Identifying the source of error also has direct implications for the precision limit of detecting change within an individual (e.g., MDC_95_). Within-subject study designs minimize the influence of between-subject baseline differences by using each person as their own reference. As such, the statistical efficiency for detecting a change, given that a change has occurred, is dominated by the residual measurement noise *σ*^2^_error_. Replicate averaging within a single session improves precision, but only up to a point: as the number of within-session measurements grows, the residual measurement noise approaches a floor set by sources of variability that are shared across all measurements within that session (e.g., fatigue state or habituation for cognitive testing, head positioning or hydration level for structural MRI, diurnal variation or time since a medication for blood-based biomarkers). Figure 5 illustrates examples of this floor for realistic noise encountered in measurement of hippocampal volume (in mm^3^; noise estimated from Elliott et al., 2026).

**Figure 5.**
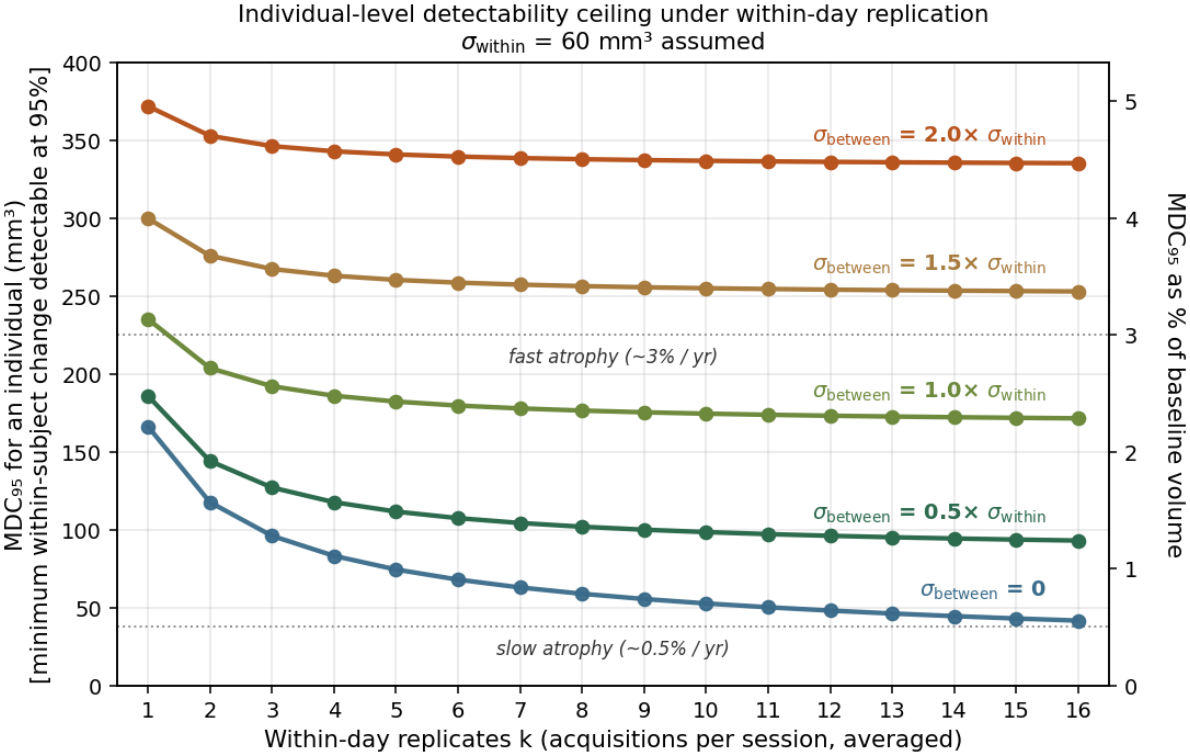
The 95% minimal detectable change (*MDC*_*95*_) for individuals as a function of *k* within-day acquisitions averaged at a single session per timepoint, at five assumed between-day noise levels (colored lines). *MDC*_*95*_ is the minimum within-subject change a single individual would need to exhibit before that change is statistically detectable. As *k* grows large, the within-day term vanishes and *MDC*_*95*_ asymptotes – a ceiling on individual-level detectability that no amount of within-session averaging can break. Curves are anchored at *σ*_within_ = 60 mm^3^, with *σ*_between_ set to multiples of *σ*_within_ spanning a realistic observed range from measurement of brain volume across MRI scans. The blue line marks the idealized *σ*_between_ = 0 reference; the four green-to-orange lines mark realistic between-day noise regimes from 0.5× to 2.0× *σ*_within_.

To push precision further, the residual error benefits from being decomposed so that within- and between-day contributions can be addressed separately. A simple modification to the duplicate change-score design makes this decomposition possible: collecting same-session replicates in addition to between-day replicates.

When measurements are replicated both within sessions and across days, *σ*^2^_error_ separates into within-session error (*σ*^2^_within-error_) and between-day error (*σ*^2^_between-error_), and each component can then be targeted by an appropriate design choice – more measures per session for the former, more days or design adjustments for the latter^7^. The relative magnitude of the two components, estimated from pilot data, determines which design choice yields the greater precision gain per unit of added measurement burden. As can be further visualized in Figure 6, precision improvement from within-session replication can reach a point of diminishing returns rapidly. Under such conditions, additional days are required to increase measurement precision.

**Figure 6.**
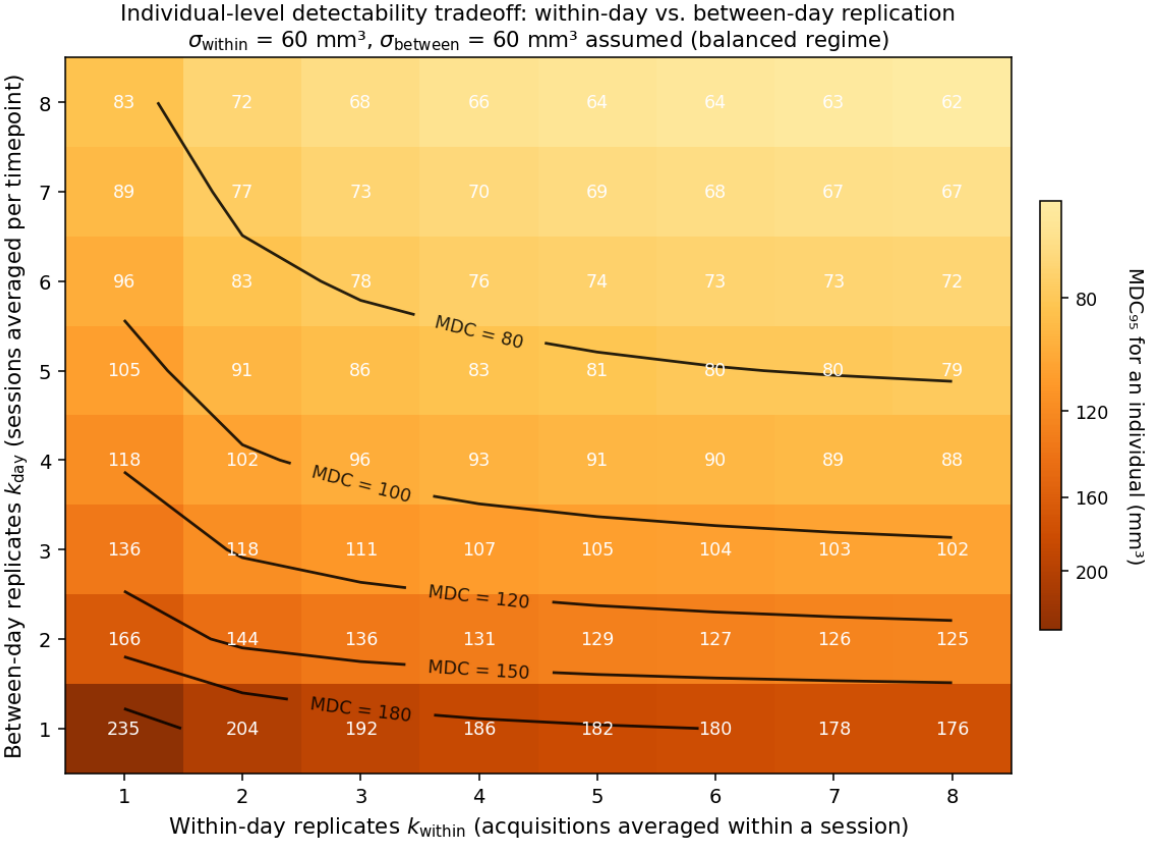
The individual-level *MDC*_*95*_ tradeoff space across combinations of within-session (*k*_*within*_, x-axis) and between-day (*k*_*day*_, y-axis) acquisitions. Each cell is labeled in mm^3^. Black contour lines mark equivalent-detectability frontiers; moving along such a frontier identifies combinations of *k*_*within*_ and *k*_*day*_ that yield the same individual-level precision. The figure anchors on the balanced regime *σ*_within_ = *σ*_between_ = 60 mm^3^ to make the asymmetry between the two axes visible. Under this regime, between-day replication is the more powerful lever – moving from a single pre+post acquisition at *k*_*within*_ = *k*_*day*_ = 1 to two between-day sessions (*k*_*day*_ = 2) reduces *MDC*_*95*_ from 235 to 166 mm^3^, whereas going to four within-day acquisitions (*k*_*within*_ = 4) on a single day reduces it only to 186 mm^3^. The picture inverts when *σ*_within_ dominates. Selection of the optimal design depends on the estimated composition of measurement error.

In the balanced regime shown, where *σ*^2^_within-error_ and *σ*^2^_between-error_ are of similar magnitude, between-day replication is the more powerful lever, because each additional acquisition reduces both noise components while additional within-session measures only reduce the within-session component. For example, reaching an MDC_95_ of approximately 100 mm^3^ requires increasing the number of days to at least three or four. Different measures will have different relative magnitudes of *σ*^2^_within-error_ and *σ*^2^_between-error_, producing distinct optimal designs. Critically, estimating these two components provides the inputs needed to allocate measurement effort efficiently between within-session and between-day replication in a contemplated future study.

## Why Have the Duplicate Change-Score and Related Designs Been Underutilized?

The logic underlying the duplicate change-score design and its variations is present across several methodological traditions, many of which emphasize that the error variance around measuring change within a person operates differently from between-subject variance, a central issue in this paper. For example, in Mellenbergh and van den Brink (1998)’s discussion of the measurement of individual change, the authors note that “a common misconception is that low reliability implies low measurement precision” (p. 471). Mellenbergh (2019) further provides paradoxical examples showing that low reliability in raw scores does not equate to poor within-subject measurement precision.

Bland and Altman’s (1986) limits of agreement methodology advocated using replicate differences to estimate measurement error and noted that high between-subject correlations can mask critical measurement properties. Specifically, their estimation of the standard deviation of change scores is similar to the present paper’s estimation of 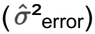 (see also Eq. 14 in Weir, 2005). Senn’s (2002) rigorous treatment of crossover designs develops the precision advantages of within-subject estimation and notes the irrelevance of between-subject variation to within-subject comparisons, but does not highlight the implications for reliability study design. Cronbach’s generalizability theory provides the formal statistical basis for the variance decomposition described here, distinguishing person variance, occasion variance, and their interaction (Cronbach et al., 1972; see also Shavelson and Webb, 1991; Brennan, 2001). Guyatt and colleagues’ (1986) N-of-1 trial methodology comes close in practice to implementing a duplicate change-score design, generating replicate change estimates within individuals through multiple treatment cycles, though its focus is on individual clinical decisions, not exploration of measurement properties.

It is important to further note that the full duplicate change-score design is only necessary in specific instances; the need to estimate change scores separately from measurement error is quite general. For example, a single change score across an intervention is sufficient to estimate effect size, which is the critical input to many design planning decisions. Test-retest raw scores at a single timepoint, such as from the baseline timepoint, alongside a single change score, provide a great deal of information about measurement properties. In its minimal form, this design requires three sessions rather than the duplicate change-score design’s four sessions. In many situations – maybe even most situations – such data would be sufficient for power planning.

That is, if a comparison’s goal is primarily to ensure signal attenuation has been avoided, and it can be assumed that the error is stable before and after the change, then collecting three sessions is likely adequate. The duplicate change-score design and its variations are most consequential when decomposition of the sources of error becomes central to experimental design and the reliability of the measures is sufficient to support such a decomposition. They are also important when there is a need to estimate the structure of the measurement error before and after the change because it cannot be assumed to be stable.

There is also a barrier to getting investigators to run designs that use replicate measures because they require increasing the number of measures per subject, which requires resources, increases subject burden, and demands multiple instrument forms and strategies to address order and item/version effects. As within-subject designs become more prevalent, the cost of not explicitly targeting within-subject precision at the design stage increases and may justify the additional burden of the duplicate change-score design. The present paper offers a practical recommendation: in a pilot or reliability study developing measures of within-subject change, investigators should consider administering pre- and post-measurements more than once per subject.

## Practical Opportunities and Considerations for Estimating Change Scores

The duplicate change-score design or its reduced three-session variant, together with its statistical formalization, is intended to provide tools to help select measures and optimize designs for within-subject research. The design is especially useful in research settings where multiple, increasingly complex, and incompletely understood derived measures index behavioral and biological constructs, and choices must be made about which processing steps to perform (for discussion see Carp, 2012; Botvinik-Nezer et al., 2020; Bhagwat et al., 2021). Once a change-score dataset has been collected, it can be reused to calibrate new processing pipelines for their power to detect within-subject change. Given the risks of overfitting (described in Appendix II), multiple datasets that allow out-of-sample testing would ideally be available as a safeguard. This section discusses challenges and opportunities related to obtaining duplicate change-score design data.

### Multiple test versions

A practical barrier to implementing the duplicate change-score design is that it requires multiple matched versions of an instrument to be available – versions that are equivalent in their measurement properties but sufficiently distinct in their surface form to minimize contamination across administrations. This requirement poses little difficulty for many biomarker assays and some neuroimaging protocols, where the measurement procedure is largely the same across occasions and the subject’s awareness of the measurement does not meaningfully contribute to its value. Order and item effects pose a greater barrier for behavioral and cognitive measures, and for brain measures that depend on task administration, where alternate forms must be carefully constructed and validated to ensure equivalence. This is a general challenge for repeat testing of behavior and, from a statistical modeling perspective, is handled well in modern theoretical frameworks and available statistical packages (e.g., Webb, Shavelson, and Haertel, 2006; Vispoel, Morris, and Kilinc, 2018).

### Tag-on measurements

It is not always practical to run the target intervention in a pilot or reliability study. For some translational programs, the question at hand is whether an intervention works at all, so the notion that power optimization depends on implementing an intervention effect creates a classic chicken-and-egg problem. Where an intervention is known to affect certain outcomes and is already being undertaken, replicate measurements can be tagged on to achieve a duplicate change-score design. For example, in an ongoing study of bipolar disorder, where there is established efficacy for mood stabilizers (e.g., lithium), but the mechanism of action is unknown, an investigator might tag on pre- and post-measurements to an existing treatment trial or an ongoing treatment clinic to learn about measurement properties of an exploratory instrument.

### Parallel interventions and mini-interventions

Another strategy is to use a *parallel intervention* or a *mini-intervention*. These are interventions that seek the same change in the measurement as the intended intervention, but that are mechanistically distinct. For example, optimizing a study design to explore treatment for major depressive disorder based on deep brain stimulation (DBS) might benefit from knowing the measurement properties of a novel behavioral assessment of rumination. Deep brain stimulation is an uncommon procedure, so a more common parallel intervention might be utilized in a pilot study to understand the measurement properties of the new instrument (e.g., cognitive behavioral therapy, standard-of-care drug treatments, or electroconvulsive therapy [ECT]). The challenge is understanding the degree to which treatment-specific mechanistic effects influence properties of the measurement in unknown ways.

Mini-interventions refer to inductions of change designed specifically as experimental tools for measurement development. They have properties of the targeted intervention but are easier to implement; the downside is that they involve clear differences and therefore require assumptions for interpretation. Building on the example above, imagine an investigator develops a behavioral instrument that measures rumination by analysis of daily voice diaries, but there are several options for how to administer the instrument. In anticipation of using the measurement instrument in a large-scale intervention study in major depressive disorder, the investigator might run a duplicate change-score design in a group of individuals who have low mood (the “worried well”) using a mood-induction paradigm involving watching sad movies. Depending on the specific measures and their properties, such a mini-intervention – which can be conducted at relatively low cost – might provide an efficient study strategy to iterate on instrument features and processing pipeline alternatives. The utility, though, will depend on whether measurement properties estimated in the worried well generalize to the clinical sample, and whether the sad-movie induction changes the measurement instrument in similar ways to how genuine therapeutic change manifests. For instance, the linguistic features that shift in the voice diaries after a sad-movie induction may or may not be the same features that shift under genuine therapeutic change in a clinical cohort. The success of such mini-interventions rests on these assumptions, which would presumably be grounded in expert knowledge about the domain and instruments, and then re-evaluated based on successes or failures in generalization.

### Surrogate interventions

When it is not possible to run an intervention in a reliability study, it may still be possible to use a well-known, controllable experimental manipulation as a stand-in (a surrogate) to mimic the signal change that is expected from the intervention. For example, in intervention studies that use task-based fMRI neuroimaging, the same task paradigm is administered pre- and post-intervention. The hypothesis is that a brain region or network will increase or decrease its signal level following the intervention in response to the consistently administered task probe. In one such study, Chuah and Chee (2008) conducted a double-blind, placebo-controlled, crossover study to test whether donepezil, an acetylcholinesterase inhibitor, could change the brain’s response during working memory in sleep-deprived subjects. The specific measurement was the fMRI response to the same working memory paradigm administered before and after the intervention.

In such a situation, a surrogate approach might be feasible to predict power and to optimize the processing stream to maximize that power. It is well established that increasing working memory load will increase the brain’s response in regions important for cognitive control (e.g., Barch et al., 2013). A surrogate duplicate change-score design can be constructed by evaluating subjects performing a low-load version of the task on one day and then a high-load version of the task on another day. This will cause the brain response to change between days, mimicking the expected intervention effect but in a reproducible, experimentally controlled manner.

### When the intervention is the passage of time

A broad class of use cases that might benefit from a duplicate change-score design focuses on change over the passage of time. In the study of child development, a central question is how the brain and cognitive processes mature, necessitating sensitive measurement of change across multiple behavioral and biological levels. In the study of advanced aging, there is intense interest in how between-subject (baseline) factors predict within-subject change in outcome measures (e.g., how does a blood biomarker measure predict brain atrophy or cognitive decline within an individual?). Longitudinal study designs can be adapted to collect duplicate change-score data by repeating assessments at each timepoint.

For example, in Elliott et al. (2026), two replicate measurement sessions were collected on different days at baseline (timepoint 1), then another pair after six months (timepoint 2), and a third pair at the one-year mark (timepoint 3). During each of these MRI sessions, a traditional brain scan was acquired as well as a burst of novel, accelerated brain scans (the exploratory measurement instrument). In such a design, it becomes possible to answer the formal question of whether the exploratory bursts of brain scans reduce measurement noise 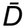 while preserving the recovered estimate of the change signal 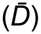 (brain aging over the year). It also becomes possible to directly estimate the power to detect change in a future clinical trial for these brain scan acquisition options as well as for alternative processing strategies applied to them (e.g., use of cross-sectional versus longitudinal morphometric pipelines, use of different software release versions). The practical takeaway is that investigators using a novel measurement instrument or processing pipeline should weigh the value of collecting multiple measurements rather than settling for just one.

## A Use Case of the Duplicate Change-Score Design Applied to Real Data

To illustrate the duplicate change-score design as a tool for instrument selection, we reanalyzed the data of Elliott et al. (2026). Thirty-eight participants were scanned with MRI across multiple timepoints, each timepoint comprising a test and retest session on different days, with the goal of tracking brain aging over one year. The measured quantity was hippocampal volume in mm^3^ extracted using FreeSurfer (Reuter et al., 2012). Four MRI acquisition alternatives were evaluated: a standard ~5-min scan from the Alzheimer’s Disease Neuroimaging Initiative (ADNI) and three novel measurement instruments (a single ~1-min compressed-sensing scan, CS1; the average of four such scans, CS4; and the average of eight such scans, CS8). Each subject contributed two change scores per instrument over the one-year interval. The scientific questions are (1) which of these four instruments would require the fewest subjects to detect hippocampal volume change in a future intervention trial, and (2) what is the required sample size for that trial? Applying the duplicate change-score variance decomposition to each instrument yields the components needed to answer these questions, summarized in Table 1.

**Table 1.**
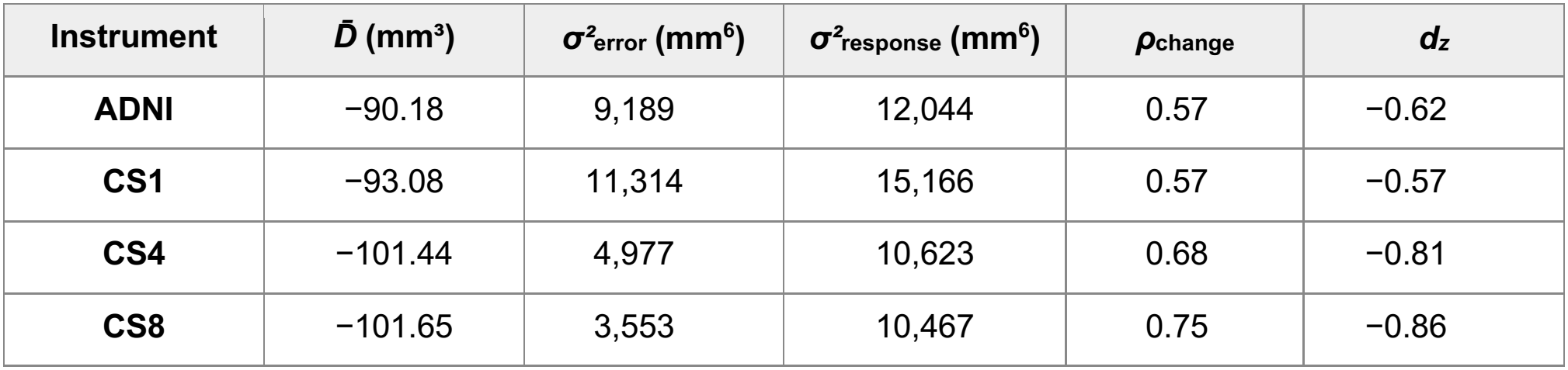
Per-instrument duplicate change-score estimates for one-year bilateral hippocampal volume change in the Elliott et al. (2026) sample (n = 38 matched subjects). Variance components are in native units (mm^3^)^2^, which is mm^6^. 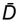 is the grand mean of the 76 change scores per instrument. *ρ*_change_ = *σ*^2^_response_ / (*σ*^2^_response_ + *σ*^2^_error_) is the change-score reliability.

| **Instrument** | ***D̄* (mm³)** | ***σ²*<sub>error</sub> (mm<sup>6</sup>)** | ***σ²*<sub>response</sub> (mm<sup>6</sup>)** | ***ρ*<sub>change</sub>** | ***d<sub>z</sub>*** |
|---|---|---|---|---|---|
| **ADNI** | −90.18 | 9,189 | 12,044 | 0.57 | −0.62 |
| **CS1** | −93.08 | 11,314 | 15,166 | 0.57 | −0.57 |
| **CS4** | −101.44 | 4,977 | 10,623 | 0.68 | −0.81 |
| **CS8** | −101.65 | 3,553 | 10,467 | 0.75 | −0.86 |

Measurement noise (*σ*^2^_error_) varies by more than a factor of three across the instruments, from 3,553 mm^6^ (CS8) to 11,314 mm^6^ (CS1). CS8 and CS4 each show significantly lower measurement noise than ADNI on a within-subject contrast (tested via a paired-bootstrap ratio; see Appendix II). Response heterogeneity (*σ*^2^_response_) is comparable across instruments (10,467 to 15,166 mm^6^), as it should be – all four measure the same change in the same people. Critically, the recovered signal 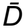 is similar for the two best-performing instruments (CS8: −101.65 mm^3^; CS4: −101.44 mm^3^), slightly reduced for ADNI (−90.18 mm^3^), and intermediate for CS1 (−93.08 mm^3^); none of these differences is significant. CS8 and CS4 are thus nominated as the more precise instruments, with no evidence that they attenuate the recovered change signal. As expected, given this pattern, change-score reliability (*ρ*_change_) rises from 0.57 for ADNI to 0.68 and 0.75 for CS4 and CS8, respectively. That is, the proportion of observed change-score variance attributable to genuine response heterogeneity grows as the measurement instrument is refined. Plotting the change scores across replicates makes the precision gains for CS8 and CS4 over ADNI readily visible (Figure 7).

**Figure 7.**
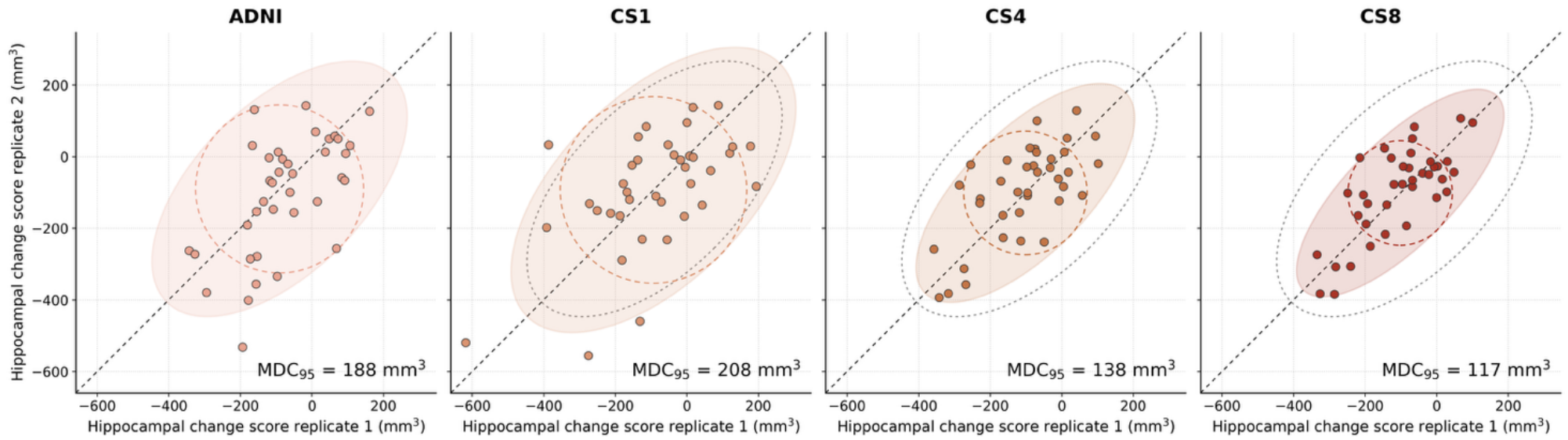
The per-instrument duplicate change scores from the Elliott et al. (2026) sample are plotted. Each point is one of the 38 subjects. The response heterogeneity, which is reflected in each subject’s position along the X = Y identity line, is separated from the estimate of the measurement error, which is the distance from the identity line (see Figure 2). The shaded ovals show the empirically estimated spread. The dotted oval in the CS1, CS4, and CS8 panels is from the ADNI reference. The dashed circle represents how measurement noise would disperse if no true response heterogeneity existed. Note that MDC_95_ is reduced for CS4 and CS8 relative to ADNI.

## Estimating sample size from the variance components.

The required sample size for a future study follows directly from the variance decomposition together with a target effect size, significance level, and power. Two decisions must be made to obtain a meaningful sample-size estimate. The first is how to anchor the expected effect 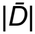. The pilot 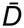 in this dataset is biologically plausible (~1.4% reduction of mean baseline volume per year), but it is also a noisy estimate at n = 38. Using it directly as the target effect for trial planning would propagate substantial pilot uncertainty in 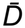 through the main trial sample-size calculation. The pilot 95% CI on 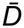 for CS8, for example, runs from −138 mm^3^ to −65 mm^3^. When squared in the power formula’s denominator, this uncertainty produces a four-fold range in plausible required-n. As a result, the required-n CI under the pilot anchor is so wide for every candidate that it has limited utility. The second decision is what *σ*^2^_response_ to assume in the future trial. The empirical pilot estimate, near 10,500 mm^6^ for the three best-performing instruments, inherits large sampling variability. Thus, a conservative sensitivity analysis should anchor 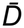 appropriately and consider a range of response-heterogeneity values

A practical approach addresses both issues by anchoring 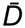 to a meaningful target change (e.g., MCID), and then sweeping *σ*^2^_response_ across the full plausible range. In many instances it will not be possible to stably estimate *σ*^2^_response_ in a sample because the reliability of the measure is not sufficient. For the present example, a target change anchor can be set at 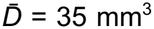, which is a change of approximately 0.50% of mean baseline and smaller than the recovered empirical effect in the pilot sample. 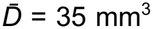 reflects the size of effect a sensitive measurement instrument would need to detect to support efficient screening of putative interventions in early-stage trials.

Required-n across a range of plausible between-subject response heterogeneity (*σ*^2^_response_) values is illustrated in Table 2. At the 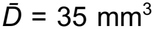 anchor, the required sample sizes shrink for the novel measurement instrument – ADNI requires about 127 subjects, CS1 about 140, CS4 about 100, and CS8 about 91 subjects. Critically, the benefit of CS8 over ADNI is present across the full range of possible response heterogeneity scenarios that might be encountered. Thus, while the actual number of subjects required to detect a change will depend on the heterogeneity of the enrolled subject sample, the benefit in power is present without exception.

**Table 2.** Required sample size to detect one-year bilateral hippocampal change at α = 0.05 (two-sided), 80% power, k = 1, for an MCID anchor of 35 mm^3^ (0.50% change from baseline) and four levels of assumed response heterogeneity. The empirical pilot *σ*^2^_response_ is approximately 10,500 mm^6^ for the three best-performing instruments. The CS8 advantage over ADNI is 36 subjects across the full sweep.

| **σ<sup>2</sup><sub>response</sub> (mm<sup>6</sup>)** | **Scenario** | **ADNI** | **CS1** | **CS4** | **CS8** |
|---|---|---|---|---|---|
| *0* | *homogeneous* | 59 | 73 | 32 | 23 |
| *5,250* | *½ × empirical* | 93 | 107 | 66 | 57 |
| *10,500* | *empirical pilot* | 127 | 140 | 100 | 91 |
| *21,000* | *2 × empirical* | 194 | 208 | 167 | 158 |

### Development of a prospective research plan

The workflow for this example provides three outputs useful for planning, summarized in Figure 8. First, the pairwise contrasts among instruments produce a ranking (CS8 ≈ CS4 > ADNI > CS1) with support about which differences are real: both CS8 and CS4 are more efficient than ADNI. Second, the variance decomposition explains the ranking: ADNI and CS1 are noise-limited (*σ*^2^_error_ about 43% of *σ*^2^_total_), CS8 is heterogeneity-limited (*σ*^2^_error_ is only 25% of *σ*^2^_total_), and CS4 falls in between. Third, the fixed-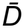 projection at the selected MCID translates the chosen instrument into a concrete sample-size estimate. At an MCID of 35 mm^3^ (0.50% change of volume from baseline per year), approximately 91 subjects are required for a heterogeneous sample. The required sample size is substantially reduced if a more homogeneous sample is enrolled. Thus, in addition to projecting realistic sample size estimates, the workflow also makes clear the benefit of subject selection.

**Figure 8.**
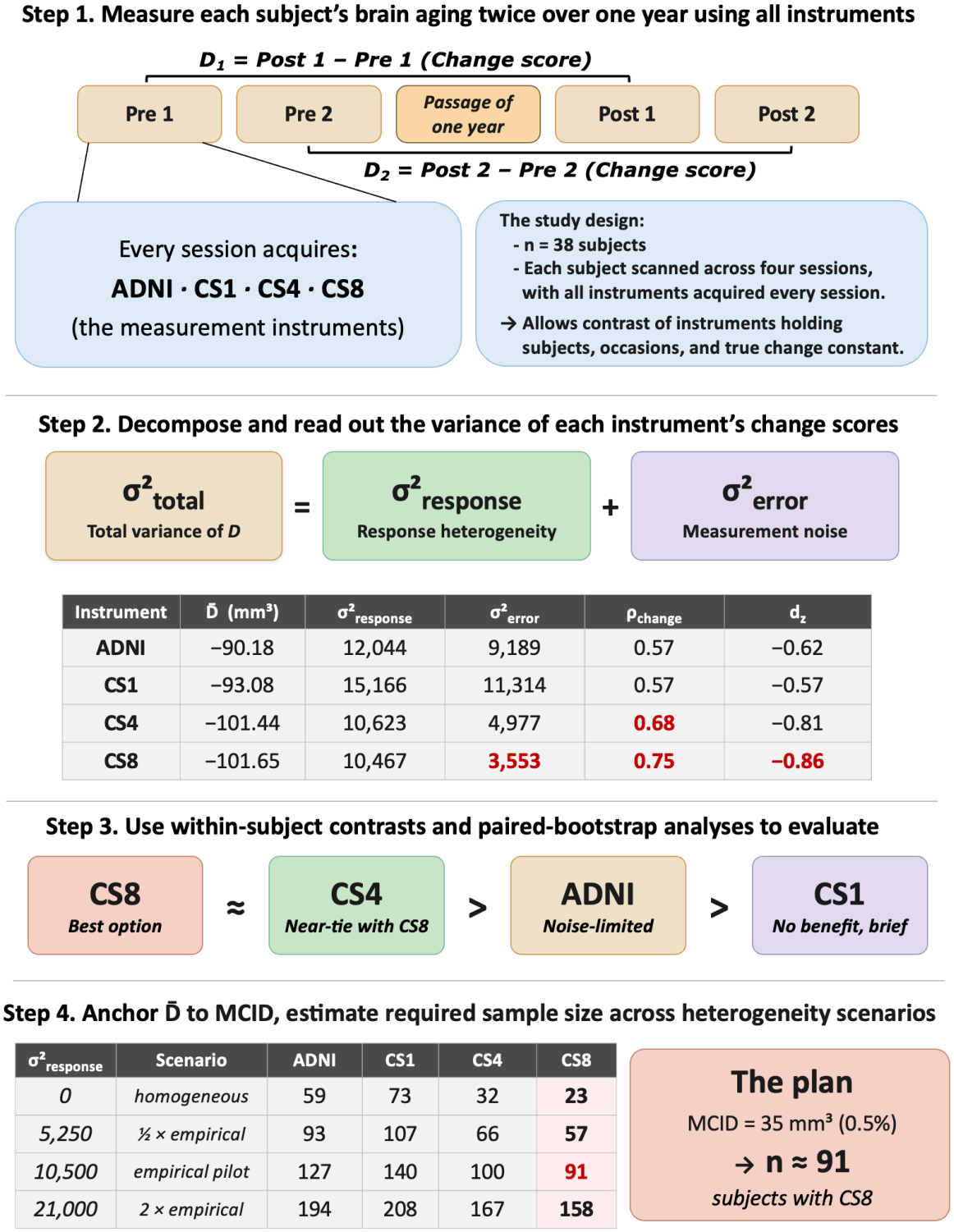
The workflow applied to data from Elliott et al. (2026) yields a power-referenced ranking and a sample-size plan. Step 1.Each subject contributes two replicate change scores per instrument over a one-year interval (*D*_*1*_ and *D*_*2*_, from the baseline and one-year session pairs), with all instruments (ADNI, CS1, CS4, CS8) acquired in every session. Holding subjects, occasions, and the underlying biological intervention constant across instruments allows for within-subject contrasts in the measurement properties themselves. **Step 2**. The variance decomposition is performed instrument by instrument, yielding 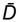, *σ*^2^_*response*_, *σ*^2^_*error*_, the change-score reliability *ρ*_*change*_, and the standardized within-subject effect *d*_*z*_. The recovered signal 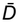 is largely flat across instruments, so the tradeoff space is one of noise reduction rather than signal attenuation. Notably, *σ*^2^_*error*_ varies by a factor of three across the instruments. Favorable estimates are highlighted in red. **Step 3**. The pairwise ranking is recovered from within-subject contrasts: paired *t*-tests on the 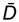 contrast (which test for signal attenuation) and paired-bootstrap ratios of *σ*^2^_*total*_ at fixed 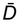 (which differ only through *σ*^2^_*error*_ and translate directly into required-*n*). The resulting ordering is CS8 ≈ CS4 > ADNI > CS1, with CS8 and CS4 both more efficient than ADNI and CS1. **Step 4**. Anchoring 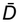 to a minimal clinically important difference (MCID) of 35 mm^3^ (≈ 0.5% of mean baseline volume) and sweeping *σ*^2^_*response*_ across the plausible range translates the estimates into a sample-size plan whose uncertainty band depends only on *σ*^2^_*error*_ sampling variability. At the empirical pilot heterogeneity, CS8 requires approximately 91 subjects to detect the MCID at *α* = 0.05 (two-sided), 80% power, against ~127 for ADNI; the absolute CS8 advantage persists across the entire sweep of response heterogeneity scenarios.

## Conclusions

By measuring twice (or more) per subject before and after a change, the duplicate change-score design estimates the mean signal change and separates measurement error from stable between-subject differences in response. The design decomposes observed variance into actionable components and, in doing so, places power planning for within-subject designs on a more solid empirical footing. A reduced design variation, where three sessions are obtained to provide one estimate of change and one estimate of measurement error, is sufficient for situations where the measurement properties can be assumed to be similar before and after the change. Expanded variations of the design further allow the decomposition of measurement error into within-session and between-day components. The design may be particularly valuable for selecting and optimizing measurement instruments in situations where numerous alternative processing pipelines are available that may attenuate the signal even as they reduce the noise. Most broadly, the framework presented here illustrates how change scores can be used to formally estimate an instrument’s measurement properties to maximize precision in within-subject study designs and to understand their limits.

## Acknowledgments and Disclosure of Generative AI Use

Generative AI was used to develop this paper. Specifically, generative AI was employed to refine the statistical arguments, explore alternatives, simulate data, draft portions of text, and proofread author paragraphs. The authors developed the idea of the within-individual repeated-measures design, its efficiency, and how its power is estimated differently from traditional use. We edited and replaced much AI-suggested text through iterations and feedback. We thank Gary King, Maxwell Elliott, and Edvard Grødem for discussion and Lin Du for checking equations. Funding was generously provided by the National Institute on Aging (R01AG067420 and U24AG092760), the Simons Foundation (811255), the Vranos Family Fund, and by Aligning Discoveries Across Psychedelic Therapies (ADAPT). ADAPT is managed by the Coalition for Aligning Science (CAS) (https://ror.org/00rf4ez40) and funded through the Sergey Brin Family Foundation (SBFF).

## Appendix I

### Efficiency of the Duplicate Change-Score Design

A concern in pilot studies is whether small samples (e.g., 30 to 50 subjects) can yield trustworthy inputs for power analyses. As described earlier, a key quantity for power analysis that is critical to within-subject design is 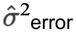, estimated as:

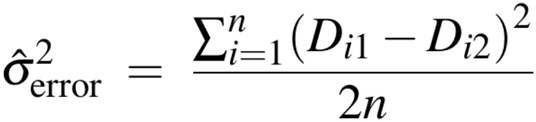

This is a mean of *n* independent squared quantities, where each squared difference (*D*_i1_ − *D*_i2_)^2^ contributes one independent piece of information about measurement error. The precision of this estimator therefore scales with n directly. The sampling distribution of 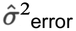 is proportional to a chi-squared distribution with n degrees of freedom:

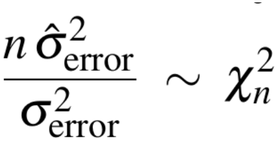

With n = 30, the 95% confidence interval around ^2^_error_ spans approximately:

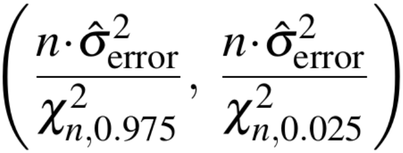

For n = 30, 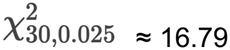 and 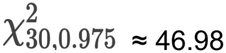, yielding a confidence interval that spans roughly a factor of 0.64 to 1.79 around the point estimate. This is sufficient precision for power planning when sensitivity analyses are conducted across a plausible range; note, however, that the confidence interval factor of 0.64 to 1.79 translates into a roughly 3× span on required sample size, so the precision is sufficient for ordering meaningfully different measurement instruments or processing pipelines but not for fine discrimination among closely matched alternatives. Increasing to *n* = 50 tightens this appreciably, but the returns beyond 50 subjects are incremental. Figure 9 illustrates how the uncertainty around the required sample size for a contemplated future study differs based on the sample size of the pilot reliability study (holding 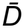 constant).

**Figure 9.**
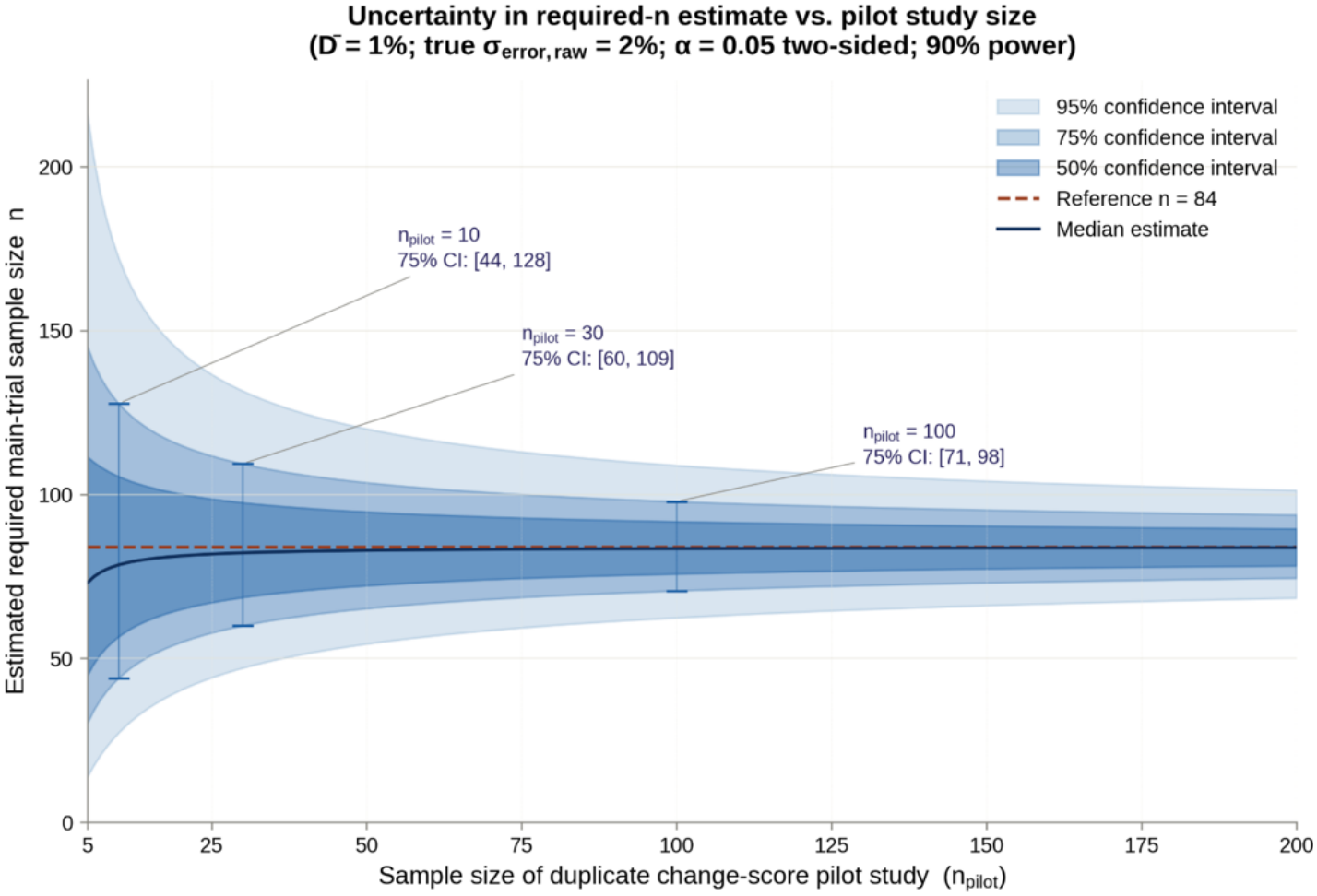
The estimated main trial sample requirements are illustrated as a function of the sample size used in the initial pilot study. This plot answers the question of what benefit investigators would get by increasing the sample size in their pilot study, assuming a fixed 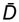. Parameters: 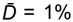, single-measurement error *σ*_error,raw_ = 2% (change-score *σ*^2^_error_ = 8 in %^2^), α = 0.05 (two-sided), and 90% power. As can be seen, the confidence interval (CI) for the needed sample size shrinks as the pilot study sample size increases. This is because the level of uncertainty is reduced. Critically, the CI shrinks rapidly and stabilizes at approximately n = 30, with diminishing returns as the sample size increases beyond about n = 50. In practice this means that a duplicate change-score design can be implemented with n = 30-50 subjects for many purposes.

Furthermore, when the duplicate change-score design is applied to compare instruments or processing streams, the propagated uncertainty of 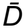 due to between-subject response heterogeneity 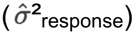 is minimized. This is because each measure is made within each subject for the same response. Given two instruments A and B, signal difference can be tested directly through a paired t-test on 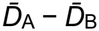. That contrast is well-estimated at modest sample sizes, even when the confidence interval around the absolute estimate of 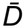 is large. If a three-session variation is used that has one test-retest replicate before or after a change, 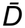 can still be estimated with slightly less precision. Appendix II describes how within-subject tests can be constructed from change-score data to compare instruments or processing pipelines.

## Appendix II

### Processing Stream Selection Can Use the Change-Score Designs as a Benchmark

Change-score designs provide candidate criteria for selecting processing stream alternatives that directly reflect within-subject measurement precision. Because each subject contributes pre- and post-measurements, the raw data can be processed through any number of competing pipelines after collection, yielding a set of change score pairs, 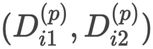, for each subject i under each processing stream p. The within-subject measurement error variance for each processing stream is then estimated as:

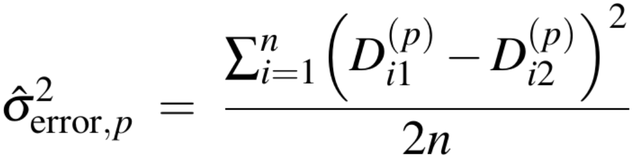

Subject-level confounds are held constant across streams because the same subjects and interventions underlie each comparison. When processing choices are being compared, the matching is even more extreme: the same raw data is used by both processing streams. Differences in ^2^_error_,_p_ across processing choices are thus attributable to the processing choice itself. The implied sample size required to achieve a target power level under each processing stream is:

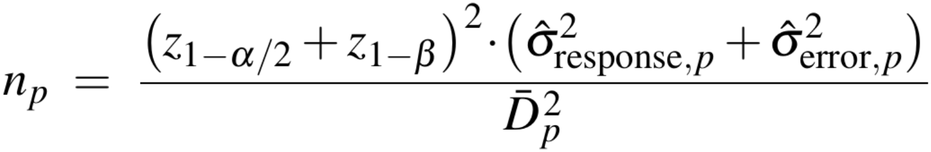

Ranking processing streams by *n*_*p*_, which is equivalent to ranking them by the standardized effect size, produces a power-referenced ordering that provides information about which pipeline requires the fewest subjects to detect the anticipated effect. Critically, ranking processing streams in this manner reflects both the effects on 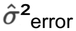 and the recovered change signal 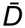. Given 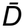 and 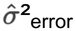 are estimated separately under each processing stream, the investigator can evaluate their independent effects on *n*_*p*_. Figure 10 shows the sample size required for two alternatives that are otherwise identical, but one attenuates the recovered signal by 30%. Figure 11 illustrates the effect on the needed sample size for a contemplated intervention trial across a broad parameter space. Note further that, for ranking processing alternatives, a single change score is sufficient to estimate the required sample size. Having duplicate change scores stabilizes the estimates (given that 2n measures are obtained).

**Figure 10.**
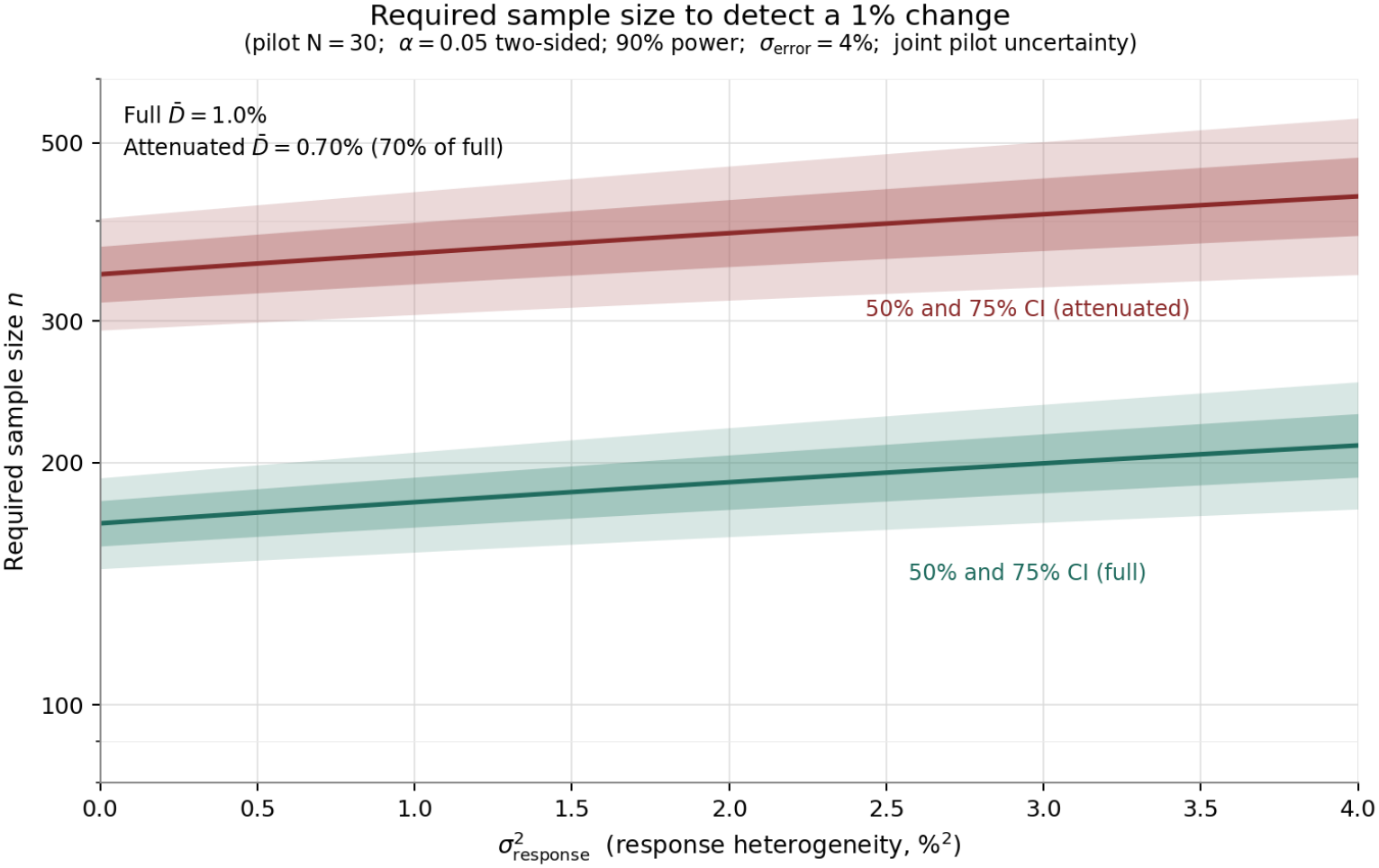
Estimated main-trial sample size requirements to detect a change as a function of true response heterogeneity, *σ*^2^_response_ (x-axis, in %^2^, i.e., the variance of subjects’ true responses expressed in units of squared percent of baseline), shown for two scenarios: a signal that is not attenuated by processing-stream choices (full 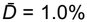 of baseline) and a signal attenuated by 30% (attenuated 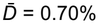 of baseline). Shaded bands show 50% and 75% confidence intervals (CI) reflecting joint uncertainty in the pilot reliability inputs (pilot n = 30). Power calculations assume α = 0.05 (two-sided), 90% power, and a fixed change-score measurement-error standard deviation of *σ*_error_ = 4% of baseline (equivalently *σ*^2^_error_ = 16 in %^2^). Sample size requirements increase substantially when the measured signal is attenuated, even when the true mean signal is held constant, and this power loss persists across the full range of response heterogeneity values examined. This plot illustrates why it is so critical to ensure that the measured signal has not been attenuated when optimizing a measurement instrument and the processing pipeline to reduce noise.

**Figure 11.**
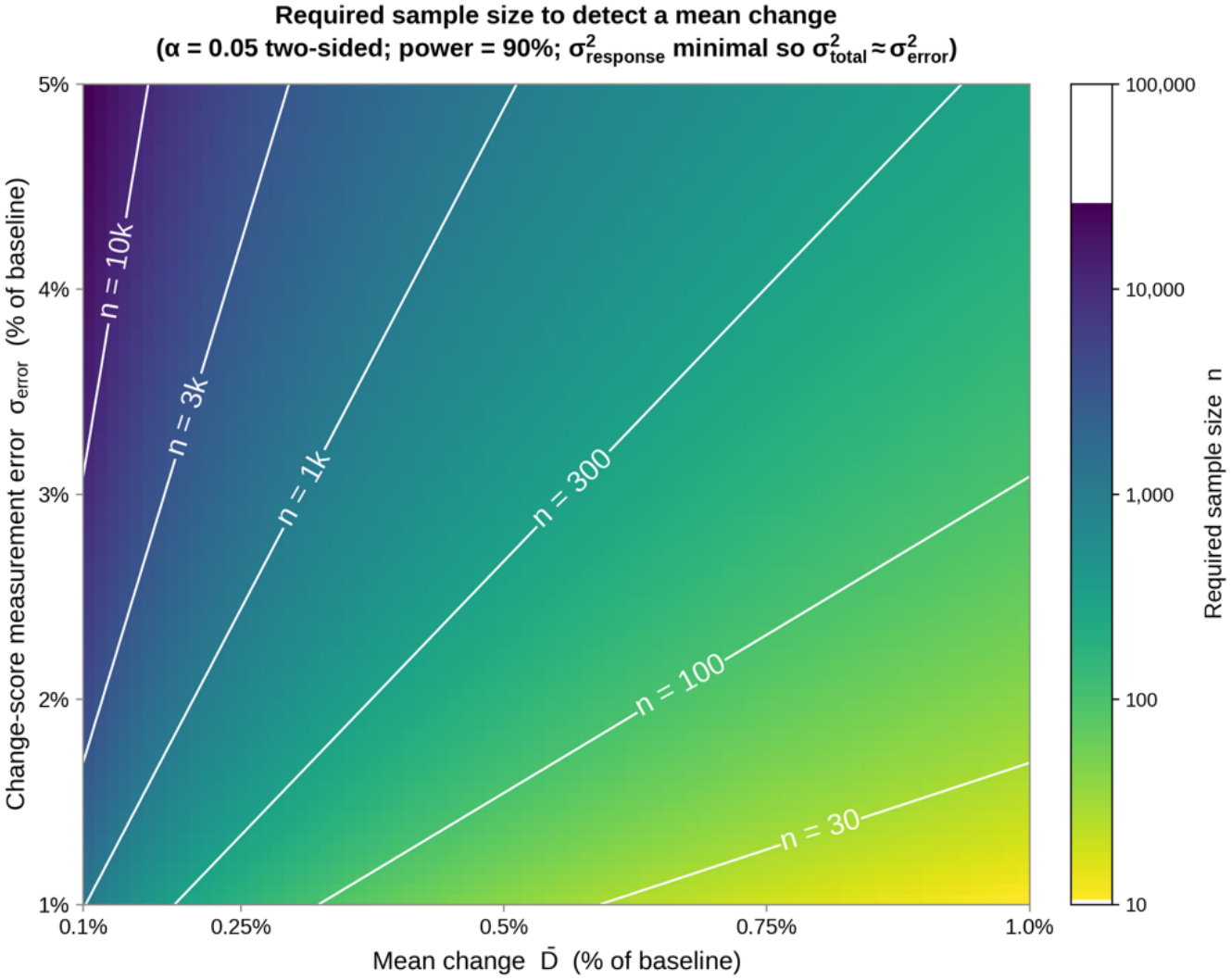
Required sample size to detect a change as a function of signal magnitude (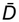, x-axis) and change-score measurement-error standard deviation (*σ*_error_, y-axis), both expressed as a percentage of baseline. The color scale shows the required sample size on a logarithmic scale; white contour lines mark sample-size isoclines (n = 30, 100, 300, 1k, 3k, 10k). Power calculations assume α = 0.05 (two-sided), 90% power, and minimal true response heterogeneity such that *σ*^2^_total_ ≈ *σ*^2^_error_. Required sample size grows steeply as the target signal magnitude shrinks or as the measurement-error standard deviation increases; modest reductions in *σ*_error_ – achievable through better instrument or processing-stream choices – can reduce sample size requirements by an order of magnitude or more across much of the parameter space.

Beyond the per-stream estimates, a change-score design can also support between-pipeline contrasts framed as paired comparisons. Because every pipeline operates on the same subjects, the between-subject 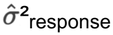 component is shared across pipelines and shifts the marginal 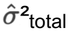 estimates by a common amount. Under these conditions, the two pipelines can be contrasted as a ratio where 1 reflects no difference, and meaningful deviations above or below 1 provide evidence for a favored pipeline. The comparison is framed as a ratio because sample-size differences change with the target effect, but the sample-size ratios stay the same – making the ratio the generalizable (scale-free) estimate of how the two instruments compare. Furthermore, signal differences (e.g., signal attenuation) can be tested separately via the paired t-test on 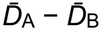. Together these within-subject tests answer the two key questions: Does this pipeline reduce noise, and does it preserve signal?

### Benchmark Decisions Using the Duplicate Change-Score Design Risk Overfitting

One risk of evaluating many processing streams against the same dataset is that the stream with the smallest estimated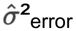 may be the one that performed best by chance in the pilot sample – a form of overfitting that would yield optimistic power projections that fail to materialize in future studies (for discussion see Simmons, Nelson, and Simonsohn, 2011; Dwork et al., 2015). Change-score designs are not immune to this challenge. This is because each estimated *σ*^2^_error,p_ combines a pipeline’s true noise properties with the chance fluctuations of the sampled subjects and occasions included in the pilot.

Selecting the minimum *σ*^2^_error,p_ from a panel of candidates in a small sample (e.g., 30 to 50 subjects) therefore preferentially identifies pipelines whose sampling fluctuations happen to be favorable in the particular dataset by chance, not only pipelines whose underlying noise properties are best. The post-selection *σ*^2^_error,p_ is, on average, a biased estimator of the value that would be obtained in a new sample, and the required-n projection that uses this same *σ*^2^_error,p_ inherits the optimistic bias. The magnitude of the bias grows with the number of candidates compared and with their similarity: when alternatives are few and their underlying error variances are well-separated, sampling noise is unlikely to change the ranking; when candidates are many and their underlying error variances are close, the winner is increasingly determined by sampling noise rather than by signal.

What addresses this bias is held-out evaluation: estimating *σ*^2^_error,p_ for the top-performing streams in a sample (or subsample) that was not used in the selection step, either through prospective replication in a new cohort or through cross-validation within the pilot sample. Pre-registering the candidate set and the selection rule before the pilot data are processed is also effective, because it bounds the implicit multiplicity that drives the bias in the first place; for decisions among well-motivated alternatives, the number of processing streams compared can often be constrained by theoretical considerations and committed to in advance.

## Footnotes

1 In contrast to *change* scores, which will be the focus of the present paper, raw-score reliability is obtained when an investigator runs a sample of subjects twice on the same instrument on different occasions and computes ICC(1,1) or a related reliability measure. The critical distinction is that raw-score ICC does not directly quantify the measurement error of within-individual change scores (but see related measures of measurement error described below), nor does it indicate whether the measure is *sensitive* to detecting change when the underlying construct truly changes between occasions – the kind of change targeted by a clinical intervention.

2 See de Vet et al. (2011, p. 112) for discussion of various definitions of SEM, including how the formulation employed here in our example can be misused when estimated from one sample and then generalized to another sample.

3 In this design variant, the intervention is administered once, and the two replicate change scores share the same underlying response. *σ*^*2*^_error_ therefore estimates measurement-instrument variability around a fixed response. A more elaborate design might repeat the intervention itself and, in doing so, capture within-subject response variability. In that case, *σ*^*2*^_error_ is inflated by intervention-to-intervention response fluctuation, and *σ*^2^_response_ reflects the stable component of response heterogeneity. Other design variations, such as having one replicate pair pre- or post-intervention (rather than at both timepoints), provide viable alternatives for many purposes. Which option is optimal will depend on the scientific question and on the assumptions that can be made.

4 An assumption of this implementation is that the first replicate change score, *D*_*1*_, does not causally influence the second, *D*_*2*_, such that the two replicates can be treated as independent observations. For measures such as hippocampal volume this assumption is reasonable – the act of measuring the construct likely does not alter future measurements of it. In many instances, particularly those involving behavioral or cognitive measures, this assumption does not hold. Practice effects, carry-over effects, and repeat-testing familiarity cause the second replicate to reflect a combination of true measurement error and systematic confounding, biasing the estimate of *σ*^2^_error_. Such situations may necessitate correction for bias, alternative models such as a mixed model with a fixed replicate / order effect and a random subjects effect, or additional experimental strategies to mitigate the early, large phase of the practice effect.

5 If one can assume that a subject’s occasion error is independent, so that their baseline replicate difference and follow-up replicate difference are uncorrelated, with no individual trend across occasions recurring, then the duplicate change-score design can also be analyzed by variance components: *σ*^2^_error_ is estimated from the 2*n* replicate pairs (the two baselines and the two follow-ups of each subject), and same-session replicates, when collected, separate *σ*^2^_within-error_ from *σ*^2^_between-error_. Under that assumption, using two error contrasts per subject rather than the single difference *D*_*i1*_ − *D*_*i2*_ estimates *σ*^2^_error_ roughly twice as precisely.

6 This is an important practical consideration: the error variance estimated in a pilot or calibration sample will generalize to the main trial to the degree that the two samples are comparable in their measurement conditions and relevant subject characteristics. For some constructs, this within-subject variability is itself a target of scientific interest rather than a nuisance to be minimized. For example, the degree of instability in repeated measurements within the same individual may serve as an informative between-subject variable in its own right – as in cases where a disease process, medication effect, or other condition systematically alters the consistency of responding.

7 Examining the sources of between-session error may indicate opportunities for increasing precision. For example, if diurnal variation is a source of measurement variation, fixing the time of testing or regressing out time of day are potential remedies. If head position within an MRI scanner is determined to be a meaningful source of occasion-specific variation, then a remedy might involve taking a person in and out of the scanner and repeating the scans.

